# Analysis of the Combined Effect of rs699 and rs5051 on Angiotensinogen Expression and Hypertension

**DOI:** 10.1101/2023.04.07.536073

**Authors:** Nicholas R. Powell, Tyler Shugg, Jacob Leighty, Matthew Martin, Rolf P. Kreutz, Michael T. Eadon, Dongbing Lai, Tao Lu, Todd C. Skaar

## Abstract

Hypertension (HTN) involves genetic variability in the renin-angiotensin system and characterizing this variability will help advance precision antihypertensive treatments. We previously reported that angiotensinogen (*AGT*) mRNA is endogenously bound by mir-122-5p and that rs699 A>G significantly decreases reporter mRNA in the functional mirSNP assay PASSPORT-seq. The *AGT* promoter variant rs5051 C>T is in linkage disequilibrium (LD) with rs699 A>G and increases *AGT* transcription. We hypothesized that the increased *AGT* by rs5051 C>T counterbalances *AGT* decrease by rs699 A>G, and when these variants occur independently, would translate to HTN-related phenotypes. The independent effect of each of these variants is understudied due to their LD, therefore, we used *in silico, in vitro, in vivo*, and retrospective clinical and biobank analyses to assess HTN and *AGT* expression phenotypes where rs699 A>G occurs independently from rs5051 C>T. *In silico*, rs699 A>G is predicted to increase mir-122-5p binding strength by 3%. Mir-eCLIP assay results show that rs699 is 40-45 nucleotides from the strongest microRNA binding site in the *AGT* mRNA. Unexpectedly, rs699 A>G increases *AGT* mRNA in a plasmid cDNA HepG2 expression model. GTEx and UK Biobank analyses demonstrate that liver *AGT* expression and HTN phenotypes were not different when rs699 A>G occurs independently from rs5051 C>T, allowing us to reject the original hypothesis. However, both GTEx and our *in vitro* experiments suggest rs699 A>G confers cell-type specific effects on *AGT* mRNA abundance. We found that rs5051 C>T and rs699 A>G significantly associate with systolic blood pressure in Black participants in the UK Biobank, demonstrating a 4-fold larger effect than in White participants. Further studies are warranted to determine if the altered antihypertensive response in Black individuals might be due to rs5051 C>T or rs699 A>G. Studies like this will help clinicians move beyond the use of race as a surrogate for genotype.

## INTRODUCTION

Hypertension (HTN) is known to involve genetic variability in the renin-angiotensin system (RAS),^1^ and one of the most implicated and studied components of the RAS is angiotensinogen (AGT). AGT is a 485 amino acid pre-pro-hormone that is cleaved by renin to the decapeptide angiotensin I, followed by angiotensin-converting enzyme (ACE) cleavage to the octapeptide angiotensin II, which causes salt and fluid retention by the kidneys to raise blood pressure. Even small increases in AGT expression have been shown to significantly increase blood pressure.^2,3^ Two single nucleotide polymorphisms (SNPs) in *AGT*, rs699 (missense SNP) and rs5051 (promoter SNP), have been associated with increased *AGT* mRNA expression and plasma AGT protein concentrations,^4-7^ suggesting a mechanistic basis to suspect these variants contribute to HTN-related phenotypes.

Rs699 A>G is a common variant that codes for a methionine to threonine (M>T) substitution at position 259 of the AGT protein (this variant has been previously referred to as M235T in the literature). An association with HTN was first published in 1992 in a study comparing hypertensive subjects to controls, and a finding of increased AGT plasma concentrations in subjects homozygous for the G allele.^7^ In attempts to find a genetic risk factor for HTN, many studies have since been conducted searching for associations between this variant and treatment response or disease susceptibility. There is controversy since some studies have not found statistically significant associations,^8-14^ but as a whole there appears to be a connection between the rs699 locus and both HTN^4,15-25^ and response to antihypertensive drugs including ACE inhibitors,^26,27^ angiotensin receptor blockers,^28-30^ and aldosterone antagonists.^31^

Of particular importance, *in vitro* evidence shows that rs699 A>G (amino acid change: M>T) has no effect on the conversion of AGT to angiotensin I by renin, which is supported by the fact that the variant does not lie within the renin binding site.^32^ Findings that rs699 A>G has no deleterious effect on protein activity suggest that disease associations could involve other mechanisms (e.g., microRNA regulation, or splice site modification) or one or more other causal variants in linkage disequilibrium (LD). The *AGT* promoter variant rs5051 C>T is in LD (r-squared 0.94) with rs699 A>G. *In vitro* assays have shown rs5051 C>T to significantly increase *AGT* transcription (up to 68.6%) through alterations in transcription factor binding.^32,33^ Due to the LD of these variants, studies in humans have not detangled the effect of each variant independently. In vitro studies of these SNPs have unraveled complex mechanistic hypotheses, but there remains a gap in understanding how each variant in isolation contributes to AGT expression in the liver (the main tissue which feeds AGT to the systemic circulation) and in important extrahepatic tissues like the kidney, brain, and vasculature.

We previously reported that (1) *AGT* mRNA is endogenously bound by mir-122-5p in human hepatocytes using the mir-eCLIP assay and (2) that rs699 A>G significantly decreases reporter mRNA levels in hepatocytes using the high-throughput functional screening assay PASSPORT-seq.^34^ Since microRNAs bind to and decrease mRNA abundance, our assay results suggest mir-122-5p decreases AGT mRNA more with the rs699 A>G variant. The putative decrease in reporter mRNA due to rs699 A>G, in theory, would oppose (i.e., “counterbalance”) the effect of rs5051 C>T on overall *AGT* mRNA expression in the human liver. The opposing effect of these tightly linked variants could make sense in the context of human evolution assuming that an imbalance in these variants would lead to unfavorable cardiovascular or other consequences. In addition to its published association with HTN, rs699 A>G has also been associated with increased power and strength performance,^35^ supporting the possibility of its role in natural selection. Both rs699 A>G and rs5051 C>T are common in humans, with the variant (rs699 G) allele frequencies ranging from 33% in Danish to 95% in African, depending on the data source.^36^ These variants occur independently from one another in about 1.3% of alleles (or 2-3% of people) based on LD-pair^37^ analysis in 1000 Genomes Project data. However, hypertension phenotypes have not yet been intentionally studied in humans where these SNPs occur independently from one another. Thus, our hypothesis is that individuals with unbalanced rs699 A>G and rs5051 C>T genotypes (i.e., having more of one variant allele than the other) would exhibit corresponding changes in AGT-related phenotypes, specifically blood pressure and the development of HTN. The objective of this study was to assess the isolated effect of rs699 A>G *in silico, in vitro, in vivo*, and retrospectively in clinical and biobank data where rs699 A>G and rs5051 C>T occur independently of each other in research participants and samples. **Table 1** provides a clear depiction of our hypothesis.

**Table 1:** Hypothesized hypertension (HTN) or *AGT* expression phenotype based on the combination of rs699 and rs5051

| Combined Genotype Group | 1 | 2 | 3 | 4 | 5 | 6 | 7 | 8 | 9 |
| --- | --- | --- | --- | --- | --- | --- | --- | --- | --- |
| Rs699 A>G | AA | AG | AA | GG | AG | AA | GG | AG | GG |
| Rs5051 C>T | TT | TT | CT | TT | CT | CC | CT | CC | CC |
| Combination Allele Frequency* | <0.0001 | 0.0063 | 0.0026 | 0.4921 | 0.4039 | 0.0818 | 0.0025 | 0.0010 | <0.0001 |
| Hypothesized HTN Phenotype (Ordinal) | ↑↑↑↑ | ↑↑↑ | ↑↑ | ↑ | ↔ | ↓ | ↓↓ | ↓↓↓ | ↓↓↓↓ |
| Hypothesized HTN Phenotype (Binned) | ↑↑↑↑ | ↑↑ |  | ↔ |  |  | ↓↓ |  | ↓↓↓↓ |

\*Alleles frequencies calculated based on all populations from the 1000 Genomes data, as accessed through the LD pair tool.

## METHODS

### In silico

We used RNAduplex to test the change in binding affinity between hsa-miR-122-5p and the *AGT* mRNA with and without the rs699 A>G variant. The ViennaRNA suite version 2.4.14^38^ was used to call RNAduplex with default settings to test the effect of rs699 A>G on binding strength. The miR-122-5p sequence was obtained online from miRbase (https://www.mirbase.org/cgi-bin/mirna_entry.pl?acc=MI0000442), and the *AGT* region of rs699 was obtained from the PASSPORT-seq oligo sequence used to clone the reporter construct.^34^ This sequence includes complementary primer ends which were included in the analysis to make sure miR-122-5p was not predicted to target these. VARNA version 3.93^39^ was used to create the RNA binding schematic figure by entering the dot bracket notation output from RNAduplex and manually editing the figure (using Adobe Illustrator version 27.1.1) to display as intended.

### In vitro

We constructed plasmid expression systems for the full-length *AGT* cDNA with or without the rs699 A>G variant on an otherwise isogenic background to test the effect of the variant on *AGT* mRNA abundance. The untagged human *AGT* cDNA expression plasmid, under CMV promoter, was obtained from Origene (catalog # SC322276) along with an empty vector to be used as a control (catalog # PS100020). The *AGT* plasmid initially contained the rs699 A>G variant (“AGT.G”) and thus was sent to GenScript for site directed mutagenesis to mutate the *AGT* cDNA from G (variant) to A (wild type, “AGT.A”). Since *AGT* is on the negative strand in the human genome, the presented nomenclature of this mutagenesis has been complemented to the positive strand. The resulting plasmids (empty vector, AGT.G, and AGT.A) were amplified in 5-alpha competent E. coli (New England BioLabs catalog C2987) and purified using Qiagen MaxiPrep according to the manufacturer’s instructions. The vector plasmid concentration was 814 ng/μL, AGT.G was 485 ng/μL, and AGT.A was 616 ng/μL, as determined by Nanodrop A260. Qubit DNA quantitation confirmed these measurements were accurate relative to each other, but Qubit-obtained concentrations were slightly higher compared to those from the Nanodrop. One million HepG2 cells were thawed from frozen stock and grown overnight in DMEM+10%FBS, resulting in 8 million cells. Cells were resuspended in 24 mL of media and 1 mL was distributed into each well of two 6-well plates. A similar procedure was used for HEK293 and HT29 cells.^40,41^ Lipofectamine 2000 was used according to manufacturer’s instructions using media without FBS to transfect 4.4 μg of plasmid per well. Cells were incubated for 3.5 hours followed by media replacement with FBS. Cells were incubated at 37 degrees Celsius for 48 hours prior to RNA isolation using Qiagen RNeasy according to the manufacturer’s protocol. TaqMan Gene Expression Assay for *AGT* (Hs01586213_m1, which does not bind near rs699) and GAPDH (Hs02786624_g1) were used according to the manufacturer’s protocol to quantify the relative mRNA abundance in cells transfected with AGT.G vs. AGT.A. Only one transfection was done for each group in the HEK293 cells, whereas three transfections were done per group in HT29, and four transfections were done per group for HepG2. Each separate transfection was considered a biological replicate. qPCR was done in triplicate for each bioreplicate. QuantStudio was used for the qPCR reactions.

To determine the proximity of rs699 to the miR-122-5p binding site, we repeated mir-eCLIP in five additional replicates of primary hepatocytes, resulting in a total of 6 mir-eCLIP datasets for analysis. The mir-eCLIP assay was performed in single-donor primary hepatocytes obtained from Xenotech (lot HC2-47) using the kit reagents and protocol provided by ECLIPSE Bioinnovations (catalog number not yet available). This protocol is developed based off the published single-end seCLIP protocol^42^ and is refined as previously described.^34^ Pooled-hepatocytes were used for Run 1 (previously completed), and data from this assay^34^ was used. Single-donor hepatocytes were used for runs 2-6. Runs 2 and 3 were performed by ECLIPSE Bionnovations and Runs 4-6 were performed by the study team. Over 50 million reads were obtained for each sample from Illumina NovaSeq 6000 sequencing. Hyb (version 1) was used to call chimeric reads from the sequencing data.^43^ A custom R pipeline was used to further analyze the Hyb output, and bedgraph files were generated for upload into UCSC genome browser.^44^ UCSC genome browser printouts were further edited in Adobe Illustrator version 27.1.1 in accordance with the UCSC genome browser user license.

### In vivo

To assess the *AGT* expression levels across genotype groups according to our hypothesis, we requested access to GTEx v8 data through dbGaP. Once approved, files were downloaded according to the AnVIL instructions provided online (https://anvilproject.org/learn/reference/gtex-v8-free-egress-instructions). BAM files were downloaded for “Liver”, “Brain - Cerebellum”, and “Colon – Sigmoid”, and featureCounts (called from the Subread package, version 2.0.1) was used to determine read counts for *AGT* for each genotype group. Rs699 and rs5051 were extracted from whole-genome sequencing VCF files using Plink version 2.0.^45^ Expression quantitative trait locis (eQTLs) and cross-tissue expression data were determined using the GTEx web browser by searching for “rs699” or “rs5051”. Web browser printouts were assembled into figures using Adobe Illustrator version 27.1.1 in accordance with the GTEx user license.

### Clinical and Biobank

To test our hypothesis in HTN phenotypes, we used 2 biobank data repositories: Indiana University Simon Comprehensive Cancer Center Advanced Precision Genomics (APG) clinic research participants,^46^ and (2) UK Biobank. APG patients provided consent for research and reporting research results generated with their data. APG data was retrospectively assessed for HTN phenotypes under a research protocol approved by Indiana University’s Institutional Review Board, utilizing available clinical sequencing data to determine rs699 and rs5051 variant genotypes for each participant. Pharmacy prescription claims data for each participant were used to determine total blood pressure medication fills per year or maximum unique blood pressure medications filled in any quarter of the year. UK Biobank data was accessed under an approved material transfer agreement and downloaded according to the UK Biobank instructions. The provided imputed genotype files were used to determine rs5051 and the measured genotype files were used to determine rs699. Plink version 2.0 was used to filter by variant and generate .raw genotype files for upload into R. UK Biobank Data fields 4080, 4079, 2966, 21022, 21000, 6177 + 6153, 131286, 21001, 20116, 22032, 1558, 1478, 30710, 30630, 30640, and 26414 + 26431 + 26421 were used to determine systolic blood pressure, diastolic blood pressure, age of HTN diagnosis, age at recruitment, ethnic background, on blood pressure medication yes/no, date HTN diagnosis, BMI (mg per meter-squared), smoking status (0=never, 1=previous, 2=current), physical activity (0=low, 1=moderate, 2=high), alcohol intake (1=daily, 2=3-4x/week, 3=1-2x/week, 4=1-3x/month, 5=special occasions, 6=never), preference for adding salt to food (1=never/rarely, 2=sometimes, 3=usually, 4=always), blood CRP, blood apolipoprotein A, blood apolipoprotein B, and education level, respectively. Sex was predefined in the .fam files provided with the UK Biobank data, which was determined by chromosome X intensity. UK Biobank genotypes were quality controlled by the UK Biobank organization (https://biobank.ndph.ox.ac.uk/ukb/refer.cgi?id=531). Further details of how the phenotype and covariate values were averaged, combined, and cleaned are provided in the R code published on github (https://github.com/Nickpowe/AGT_rs699_HTN). For example, systolic blood pressure was averaged across 1-4 visit instances for each individual. Race and ethnicity was broadly grouped into White, Black, Indian, Asian, White and Black, White and Asian, Prefer not to Answer, Mixed, and Other based on the more granular data provided by UK Biobank survey responses. Blood biochemistry data used just the first instance (whenever multiple instances existed) since most of the blood pressure values were recorded from the first instance.

### Statistical Methods

qPCR was analyzed using two-sample two-sided t-tests for data that (1) averaged the technical replicates or (2) kept technical replicates separate (both results provided). Fold-changes were calculated by comparing to the wild type group (AGT.A) after normalizing the wild type cycle thresholds to the average of all wild type cycle thresholds. All cycle threshold values were also normalized to GAPDH. GTEx eQTLs were precalculated (https://gtexportal.org/home/methods) based on normalized expression slope (slope of regression estimate) and are displayed with 95% confidence intervals. Statistical analyses of the GTEx genotypes and the APG data were not conducted due to low sample sizes in high and low genotype groups. UK Biobank data were analyzed using base R linear or logistic regression. Covariates were included as independent factors along with genotype groupings in multiple regression models. Tests for normality were done visually given the large amount of data. Rs5051 was imputed, and imputation dosages were converted to best guess by rounding to the nearest whole genotype number (0, 1, 2) when used in genotype groupings; otherwise, the imputation dosage was used in the regression. Statistical analyses were stratified into subgroups based on age <50 or ≥50, on blood pressure med yes/no, and into race and ethnicity groups. Regression estimates for systolic blood pressure, diastolic blood pressure, mean blood pressure (average of systolic + diastolic), and age of HTN diagnosis were compared by dividing the estimates by the standard deviation for the whole cohort (for each phenotype, respectively), resulting in z-score-like values. R version 4.1.1 was used for all statistical analyses and to generate all figures.

## RESULTS

### In Silico Analysis of MicroRNA Binding Site Creation

We previously reported that *AGT* mRNA is endogenously bound by mir-122-5p in liver cells using the mir-eCLIP assay and that rs699 A>G significantly decreases reporter mRNA levels in liver cells using the high-throughput functional screening assay PASSPORT-seq.^34^ Thus, we further investigated this role of mir-122-5p on *AGT* expression using *in silico* tools. The existing microRNA binding and mirSNP prediction tools (mirDB, miRdSNP, PolymiRTS, and others) focus on 3’UTRs of target genes and thus fail to consider the rs699 site, which is in exon 2 of *AGT*. Therefore, we used RNAduplex to test the hypothesis that rs699 A>G creates a stronger binding site for mir-122-5p, as this could explain the downregulation seen in the reporter assay. The results of the RNAduplex binding analysis show that the variant construct (**Figure 1**, right) predicted an additional 4 bases to pair (3 of which are G-U pairs) compared to the reference construct (**Figure 1**, left) and increased the binding strength from a deltaG of -11.3 to -12.3 kcal/mol (a lower free energy state supporting stronger binding). To put this deltaG change in context, we randomly scrambled the target sequence 2000 times and recorded the smallest deltaG and subsequently recorded the deltaG of a perfectly matching siRNA to generate the highest and lowest plausible values (−7.1 to -39.3). The largest plausible RNAduplex change was 32.2 kcal/mol, and compared to these extremes, the rs699 A>G variant increased the binding strength by 3%.

**Figure 1:**
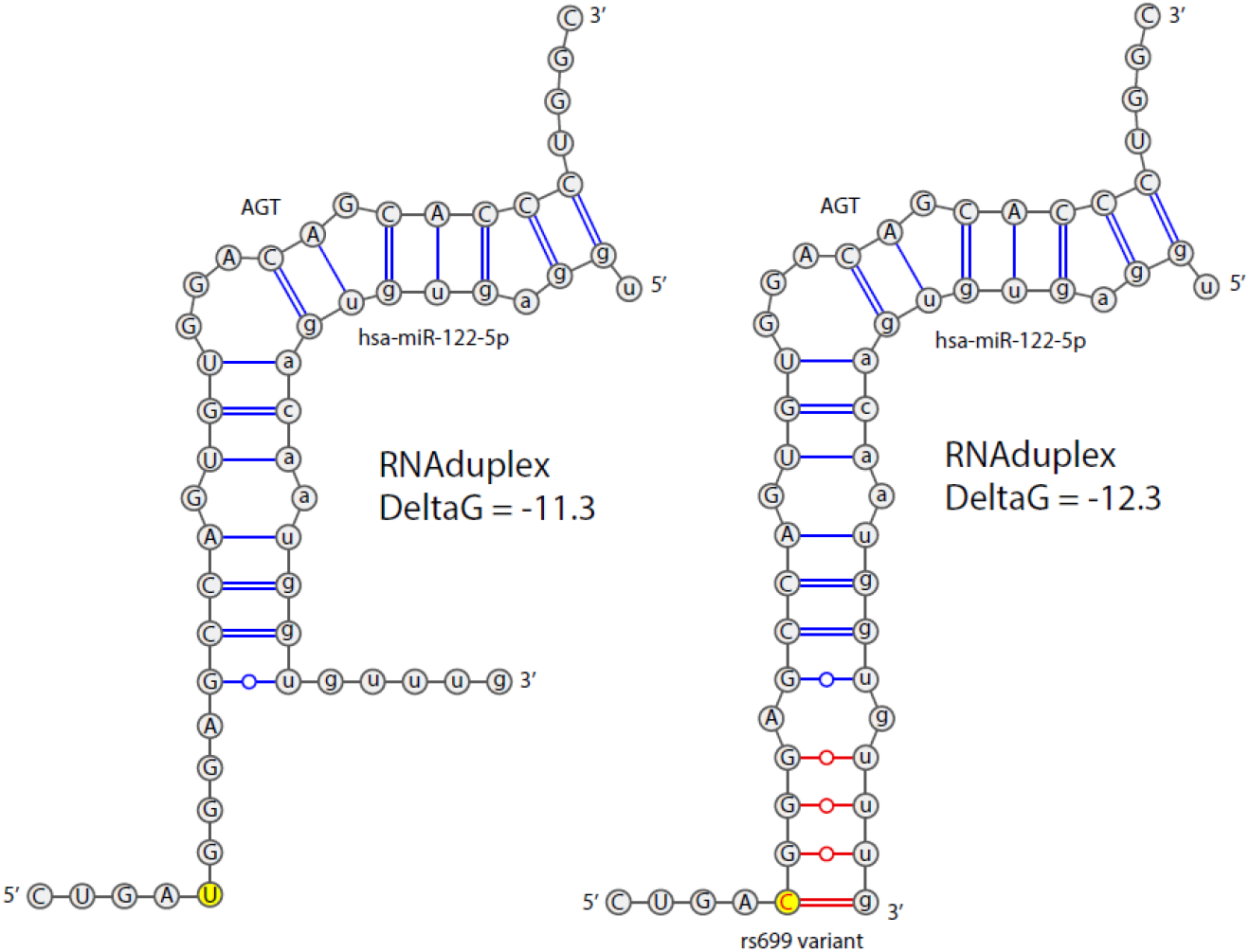
Schematic showing the RNAduplex predicted increase in binding strength with the rs699 A>G variant. The variant position is highlighted in yellow. Nucleotide bonds that are predicted to arise due to the variant are shown in red. GC pairs are shown with double lines, AU pairs are shown with single lines, and GU pairs are shown with single lines with open dots. *Note: The variant nomenclature “rs699 A>G” is complemented to the positive strand, whereas the nucleotides shown are for the AGT mRNA structure, which is transcribed from the negative DNA strand*. Abbreviations: AGT = angiotensinogen; hsa = homo sapien; miR = microRNA; DeltaG = Kcal/mol change in free energy; C = cytosine; G = guanine; A = adenine; U = uracil.

### Mir-eClip to Verify mir-122-5p Binding Sites

To further test the hypothesis that mir-122-5p binds near the location of rs699, we repeated the mir-eCLIP assay in 5 replicates on individual donor hepatocytes to gain repeated measures of where microRNAs bind in this location. We found that the rs699 location is about 40-45 nucleotides away from the strongest microRNA binding peak (**Figure 2**) in the entire *AGT* mRNA. mir-122-5p constitutes the majority of the mir-eCLIP reads, which is not surprising given mir-122-5p makes up close to 70% of the abundance of all microRNAs in hepatocytes.^34^ We also detected mir-26a, mir-26b, and several other microRNAs contributing to the main peak shown in **Figure 2**, but these were not highly prominent. This analysis demonstrates that there is microRNA binding activity occurring in the specific location of rs699, but that the rs699 A>G variant may be exerting more of its effect indirectly given the close proximity (but not exact overlap) with a very strong site of microRNA binding.

**Figure 2:**
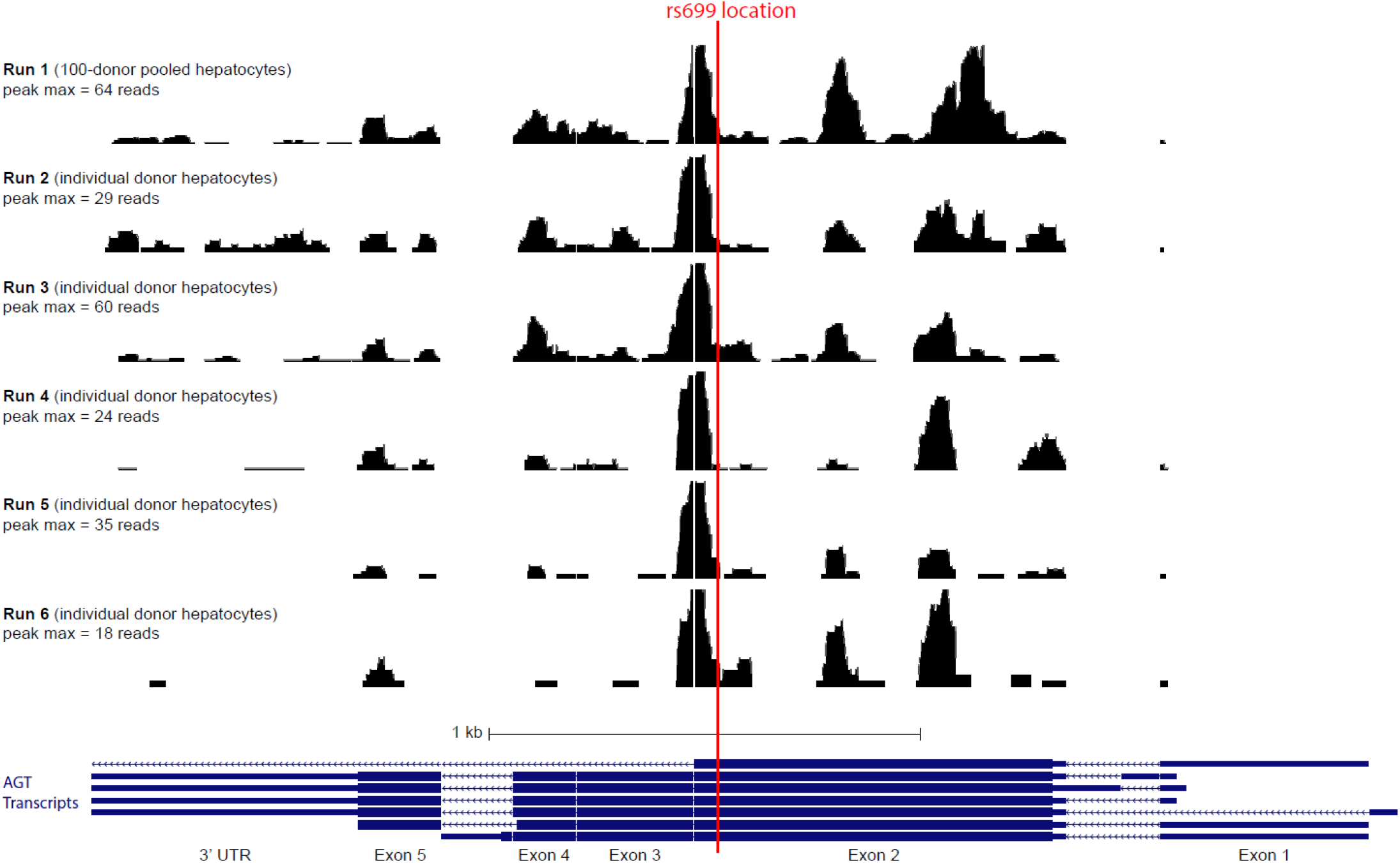
UCSC Genome Browser view of our miR-eCLIP data in BedGraph format, showing the proximity of rs699 to the most prominent microRNA binding peak in the *AGT* mRNA. Abbreviations: AGT = angiotensinogen; kb = kilobase; UTR = untranslated region; max = maximum.

### eQTL Analysis of rs699 in GTEx

Since the reporter assay demonstrated a 5-fold decrease in mRNA, and the role of mir-122-5p in regulating *AGT* mRNA abundance is supported by the *in silico* evidence and the *in vitro* mir-eCLIP evidence, we used the GTEx data to test the association between rs699 A>G and *AGT* mRNA expression in human liver. **Figure 3 (right)** shows that there is no significant change in *AGT* mRNA abundance according to rs699 genotype in the liver. However, in other tissues like the sigmoid colon and cerebellum, there is a strong significant increase and decrease in *AGT*, respectively. **Figure 3 (left)** shows the bulk mRNA expression for each tissue and reveals a possible trend towards more rs699 A>G downregulation effect in tissues expressing higher levels of *AGT*, like the brain and arteries. As expected, due to the significant LD, rs5051 C>T showed very similar results and thus these two variants need to be decoupled to address our hypothesis.

**Figure 3:**
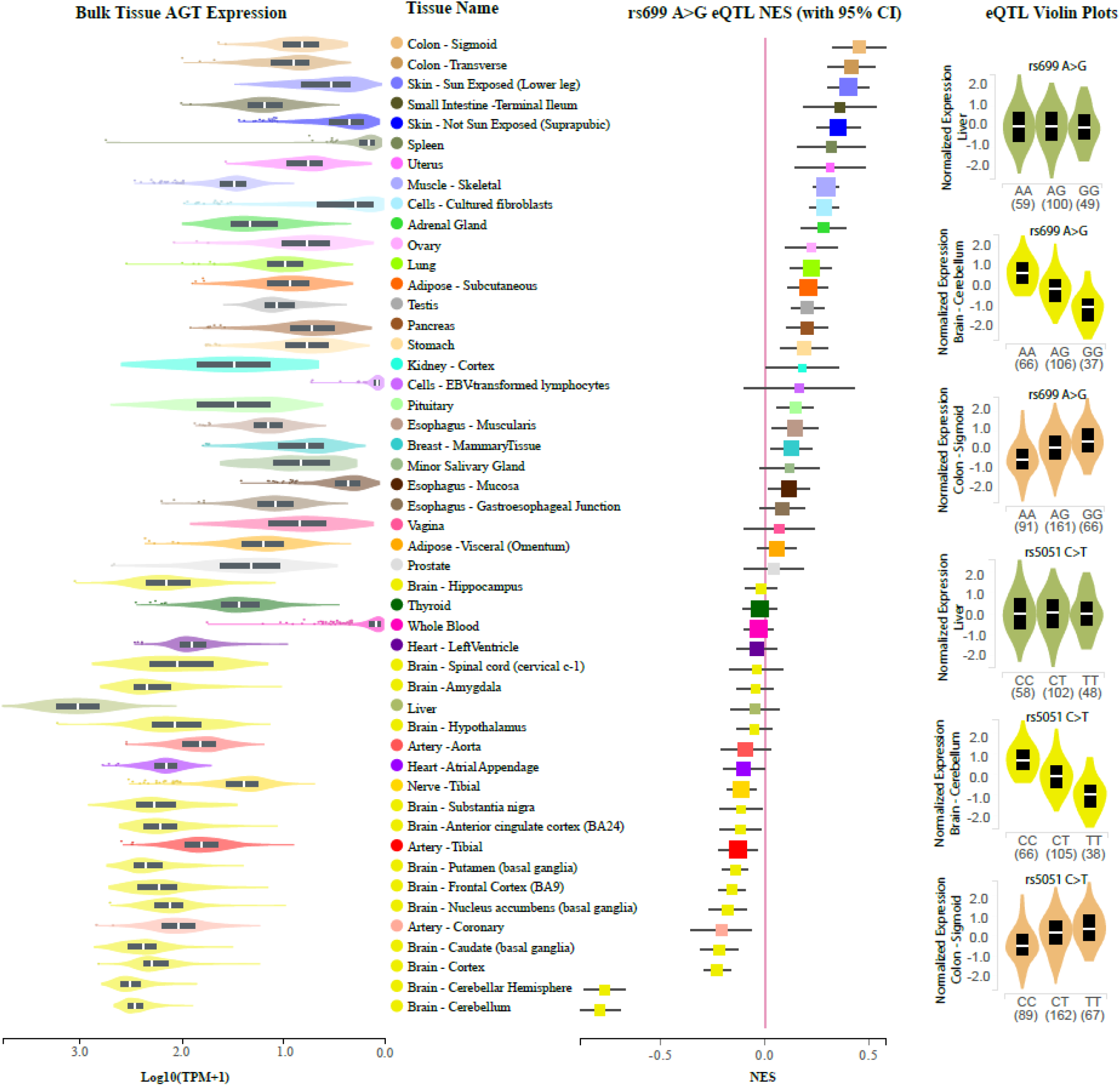
GTEx web-portal data. (Left) Bulk mRNA expression data in log transformed units. (Middle) eQTLs for rs699. (Right) Violin plots showing the more granular eQTL data for rs699 and rs5051 for Liver, and tissues with the most extreme eQTLs for rs699. Abbreviations: AGT = angiotensinogen; eQTL = expression quantitative trait loci; NES = normalized expression slope; TPM = transcripts per million.

We hypothesized the effect of rs699 and rs5051 genotypes would occur according to **Table 1**, where rs699 A>G decreases AGT expression, reduces blood pressure, and is protective of HTN, and where rs5051 C>T increases AGT expression, increases blood pressure, and contributes to increased development of HTN. In the GTEx liver data, there were three individuals who had one more rs5051 T allele than rs699 G, and three individuals who had one more rs699 G allele than rs5051 T. We tested our hypothesis according to the genotype groups described in **Table 1**. We expected to see elevated *AGT* expression in genotype groups 1-3 (more rs5051 T alleles than rs699 G) and decreased *AGT* expression in groups 7-9 (more rs699 G alleles than rs5051 T). Due to the low sample size, we could not conclude from this analysis if the two SNPs have opposing effects on *AGT* expression because there were too few numbers in the high and low groups (**Figure 4**). The analysis in cerebellum and sigmoid colon for genotype groups 4-6 demonstrated a similar trend to that observed in the individual SNP GTEx data shown in **Figure 3**, and the lower and upper genotype groups did not reveal marked effects on *AGT* expression.

**Figure 4:**
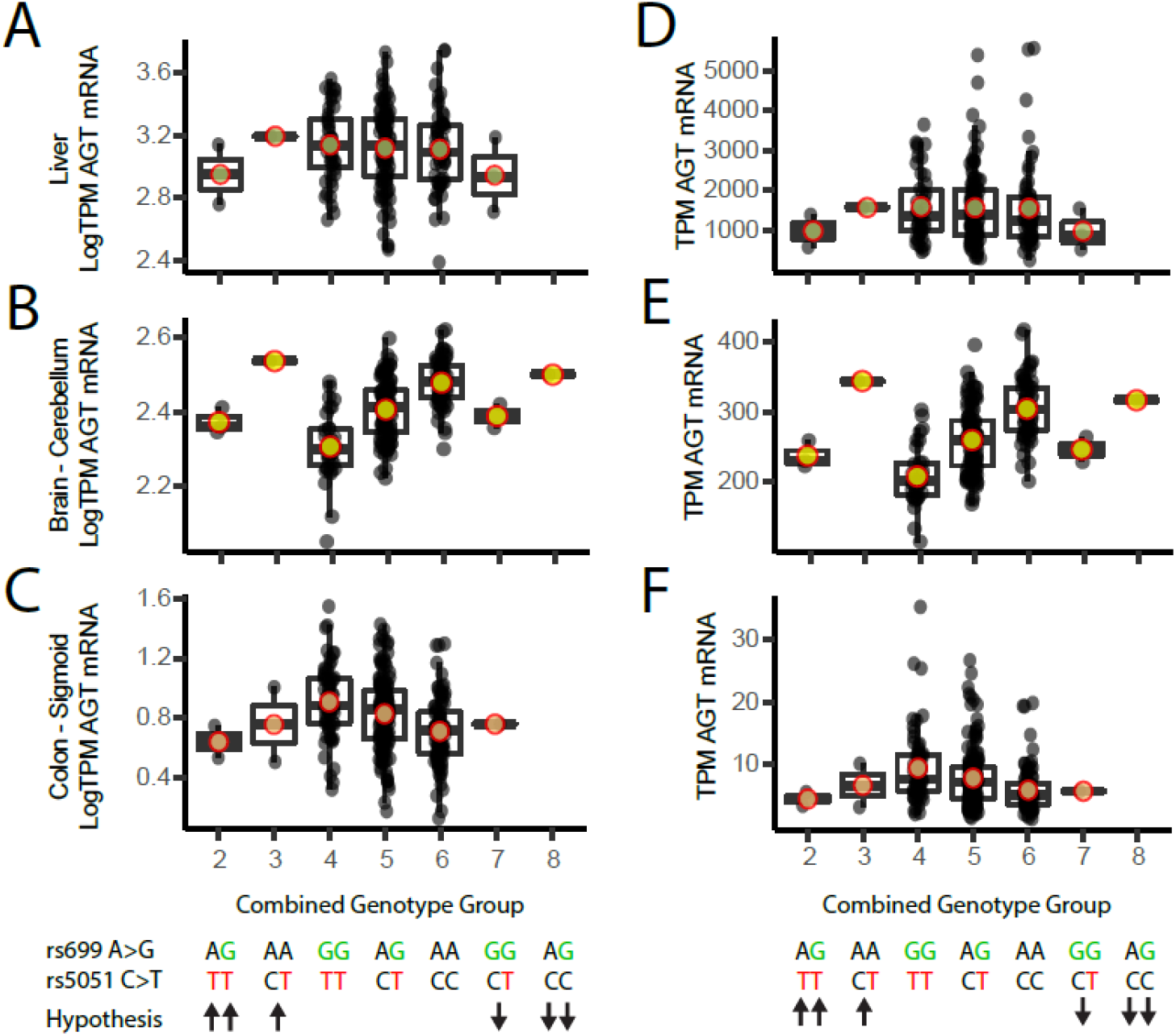
Liver *AGT* expression across the genotype groups shown with the hypothesized effect on *AGT* expression. (Left column; A-C) Log transformed TPM liver *AGT* reads. (Right column; D-F) Linear TPM liver *AGT* reads. (Top row; A, D) Liver *AGT* reads, n=208. (Middle row; B, E) Brain-Cerebellum *AGT* reads, n=209. (Bottom row; C, F) Colon-Sigmoid *AGT* reads, n=318. Abbreviations: TPM = transcripts per million, AGT = angiotensinogen.

### Mechanism in Different Cell Types

Given that significant *AGT* expression eQTLs existed in opposite directions in colon and cerebellum, we conducted transient transfections of *AGT* in three cell types to test the effect of rs699 A>G on mRNA abundance/expression from a plasmid containing the full *AGT* gene with or without rs699 A>G on an otherwise isogenic background. We could not identify a suitable cell model for cerebellum, therefore we only utilized liver (HepG2) and colon (HT29) models. We also included kidney cells (HEK293). We expected to see that in HepG2 cells, where mir-122-5p is the 48^th^ most highly expressed microRNA (out of 680 total), *AGT* would be decreased by the rs699 A>G variant. However, rs699 A>G caused a near-significant 1.8-fold increase in *AGT* mRNA of HepG2 liver cells after a 48-hour transient transfection (**Figure 5A**).

**Figure 5:**
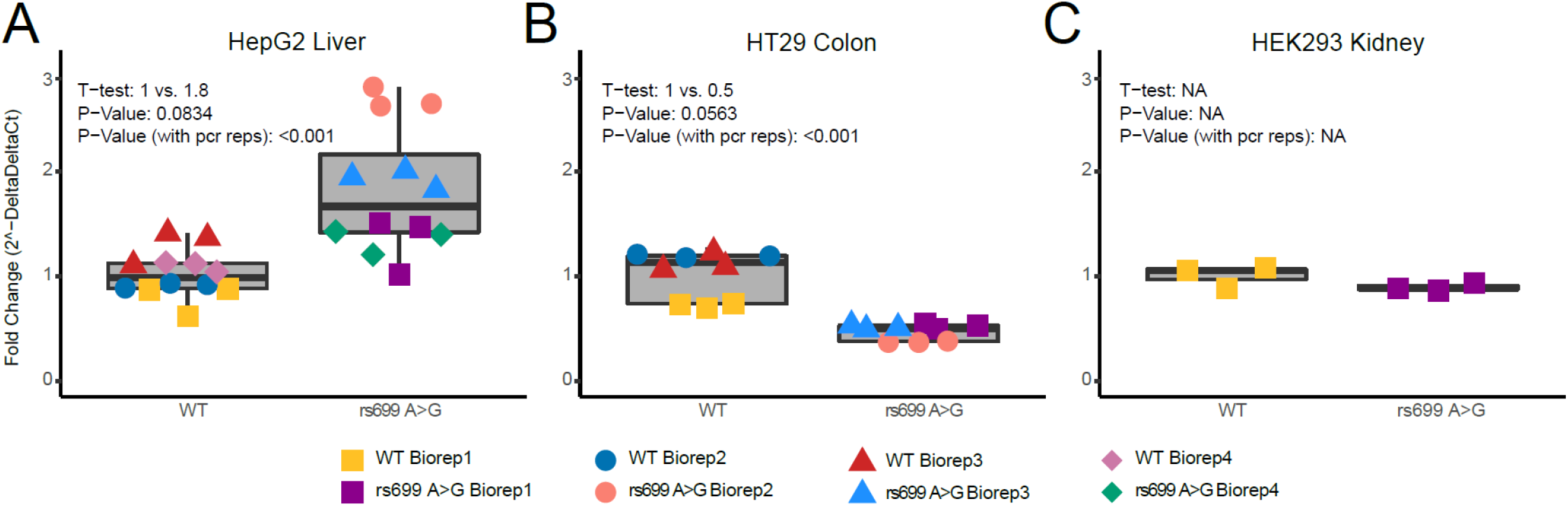
(A-C) qPCR results for 3 different cell lines. (A) *AGT* expression in HepG2 liver cells, (B) *AGT* expression in HT29 colon cells, (C) *AGT* expression in HEK293 kidney cells. Abbreviations: Ct = cycle threshold; PCR = polymerase chain reaction; reps = replicates; WT = wild type; Biorep = biological replicate; NA = not assessed. APG = advanced precision genomics

Based on the GTEx results, we expected to see HT29 colon cells express more *AGT* in the rs699 A>G construct, yet rs699 A>G caused a near-significant 2-fold decrease in *AGT* mRNA of HT29 colon cells after a 48-hour transient transfection (**Figure 5B**). Rs699 A>G did not cause a significant change (1.1-fold) in *AGT* mRNA of HEK293 kidney cells after a 48-hour transient transfection (**Figure 5C**). These results isolate the effect of rs699 A>G and indicate that cell-type specific factors are leading to seemingly differential regulation of *AGT* mRNA abundance. Given the differing effect of this SNP in different cell types, further work is needed, if warranted, to understand how *AGT* expression is modulated by this SNP.

### Analysis in Biobank data

To further test our hypothesis, and to determine if there is value in more rigorous studies of these variants, we used available clinical cohort and biobank data. The first cohort consists of cancer patients from the Indiana University Advanced Precision Genomics (APG) Clinic where we have HTN medication fill data. We divided the subjects into genotype groups as shown in **Table 1**, and **Figure 6A** and **6B** shows the results of this analysis for two phenotypes: (1) total blood pressure medication fills per year, and (2) total maximum concomitant unique blood pressure medications filled in any quarter of a year. We did not perform statistical comparisons on this data since there are too few numbers in the high/low risk genotype groups. However, it appears there could be a difference in HTN phenotypes across the genotype groups.

**Figure 6:**
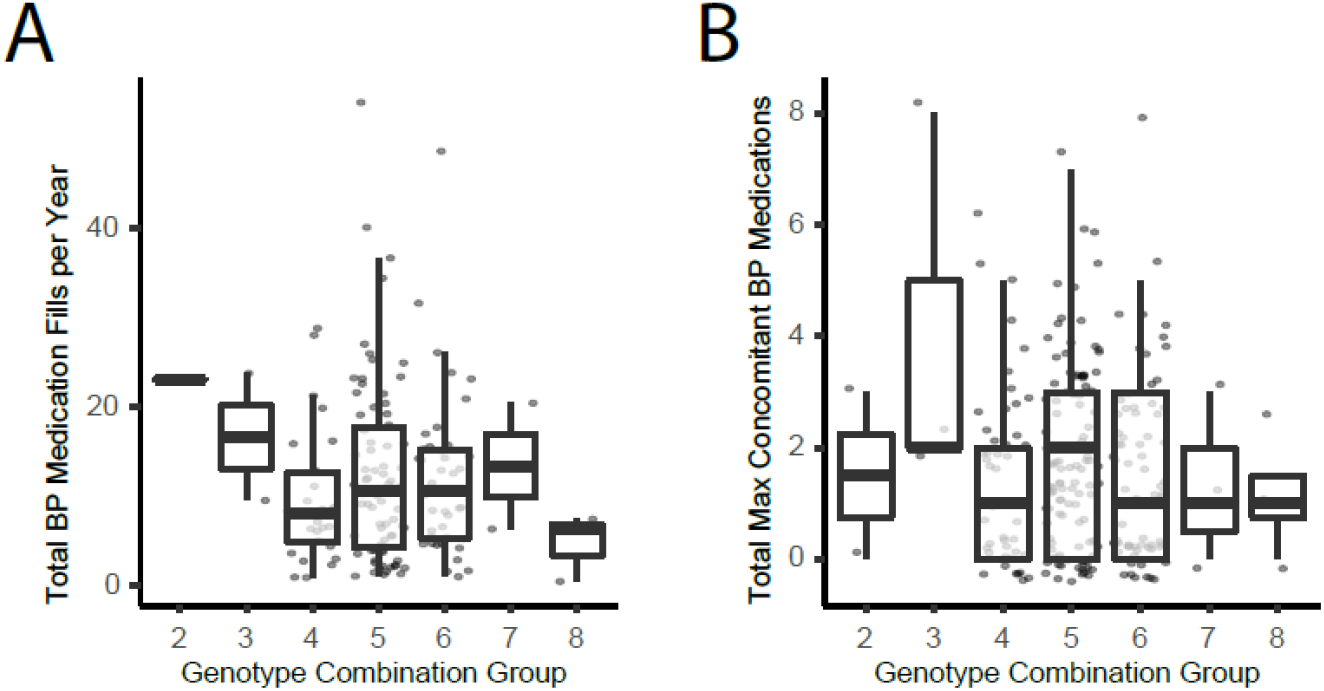
APG cohort results for (D) total fills per year, (E) total maximum concomitant meds at any given time.

The trend towards lower HTN medication usage in the groups with more rs699 G alleles (groups 7-9) appeared supportive of our hypothesis, therefore we tested our hypothesis again in a much larger retrospective analysis in the UK Biobank data.

Among 462,417 individuals tested in the UK Biobank, we found that rs699 A>G was significantly associated with a 0.11 mmHg increase in systolic blood pressure per rs699 G allele (p=0.005). Not surprisingly, rs5051 C>T was also significantly associated with a 0.10 mmHg increase in systolic blood pressure (p=0.011) in the same individuals. The direction of effect of these results are consistent with the literature when the two variants are not considered as a combined genotype. We repeated this analysis after controlling for the following covariates: sex, age, body mass index (BMI), smoking status, physical activity level, alcohol intake, preference for adding salt to food, blood c-reactive protein, blood apolipoprotein A, blood apolipoprotein B, and level of education. These were chosen because they have been previously found to correlate with or be predictive of HTN.^47^ After controlling for these factors, we found that rs699 A>G was significantly associated with a 0.36 mmHg increase in systolic blood pressure per rs699 G allele among 311,004 individuals (p<0.001). Not surprisingly, rs5051 C>T was also significantly associated with a 0.35 mmHg increase in systolic blood pressure (p<0.001) in the same individuals (**Figure 7C** and **7F**) per genotype group.

**Figure 7:**
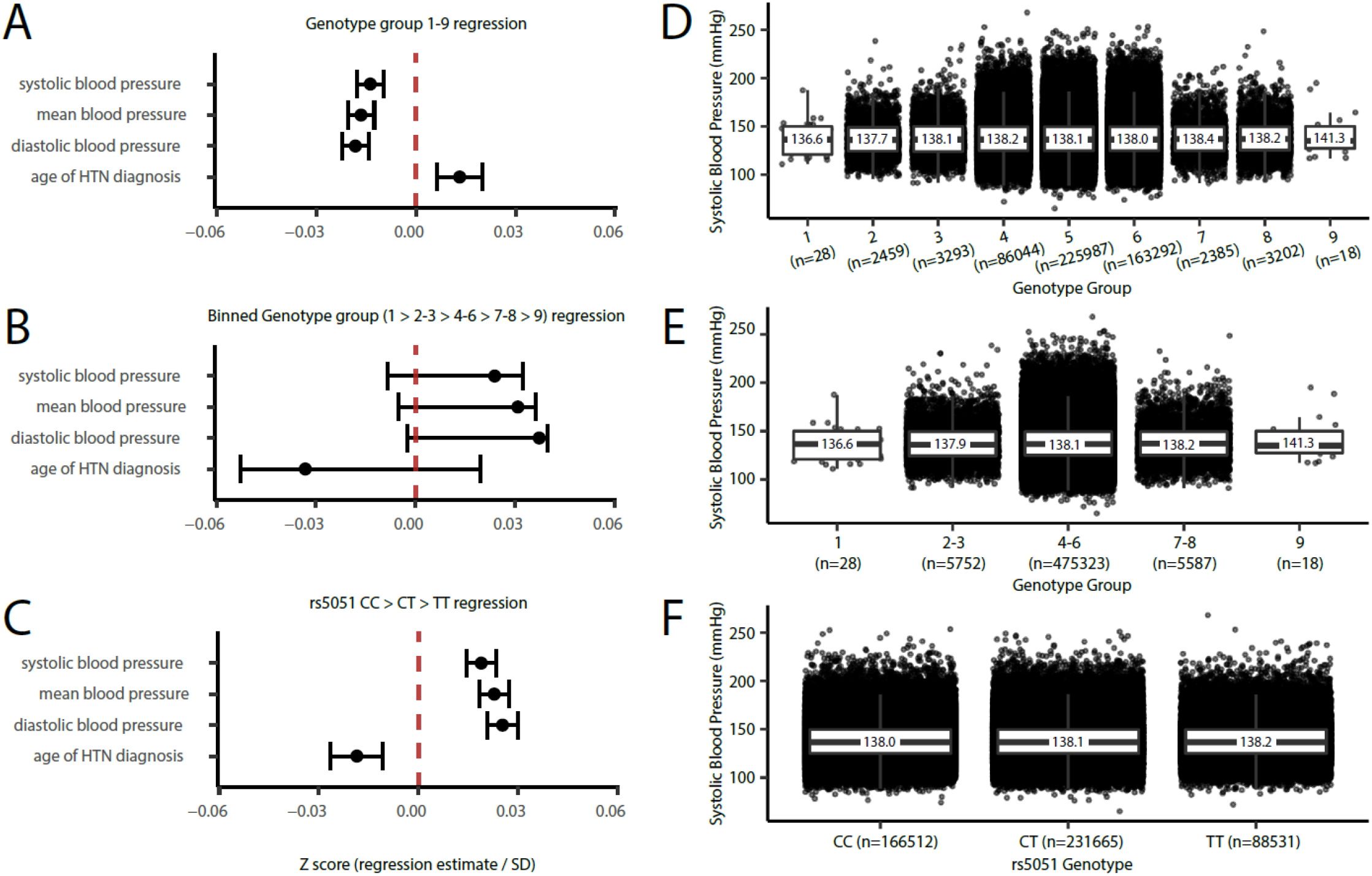
Blood pressure phenotype regression estimates (corrected for covariates and normalized to standard deviation) with (A) genotype groups according to our ordinal hypothesis, (B) binned genotype groups that test only groups with unbalanced rs699 G and rs5051 T, and (C) rs5051 genotype group. Systolic blood pressure boxplots are shown for (D) genotype groups according to our hypothesis, (E) binned genotype groups that test only groups with unbalanced rs699 G and rs5051 T, and (F) rs5051 genotype group. Boxplots display median (black bar), 25^th^ to 75^th^ percentile (box), and mean values are labeled over the box.

When we conducted the analysis by including rs699 and rs5051 genotype combinations according to our hypothesis, and controlling for the same covariates, there was a statistically significant effect that supported our hypothesis that an imbalance favoring rs699 G over rs5051 T (i.e., genotype groups 7-9) would decrease systolic blood pressure (linear regression estimate= -0.25 mmHg change per genotype group, p<0.001). **Figure 7A** shows these results as Z scores (normalized to the standard deviation of the full cohort) along with the similar results for mean blood pressure, diastolic blood pressure, and age of HTN diagnosis. However, when we inspected the blood pressure averages for each genotype group (**Figure 7D**), it appeared that the regression was being driven by the large sample size in genotype groups 4, 5, and 6, and that the lower and higher genotype groups (1-3 and 7-9) did not follow the hypothesized pattern. To further isolate the effect of rs699 A>G, we binned genotype groups into 1 vs. 2-3 vs. 4-6 vs. 7-8 vs. 9, and found that there was a non-significant trend towards an increase in blood pressure (**Figure 7B** and **7E**) which statistically demonstrates that rs699 A>G is unlikely to be exerting a blood pressure lowering effect. **Table 2** provides the regression estimates for the systolic blood pressure phenotypes, including estimates for the covariates.

**Table 2:** Regression results for systolic blood pressure in the UK Biobank. Variable ranges are described in the methods.

|  | Genotype group 1-9 |  | Binned genotype group |  | Rs5051 |  |
| --- | --- | --- | --- | --- | --- | --- |
|  | Estimate | p-value | Estimate | p-value | Estimate | p-value |
| (Intercept) | 73.82541 | <2.22E-308 | 71.86081 | <2.22E-308 | 72.21228 | <2.22E-308 |
| Group | -0.25438 | 8.51E-12 | 0.220297 | 0.252838 | 0.34963 | 1.51E-16 |
| SEX | -6.45016 | <2.22E-308 | -6.45415 | <2.22E-308 | -6.44912 | <2.22E-308 |
| age | 0.671527 | <2.22E-308 | 0.671052 | <2.22E-308 | 0.671726 | <2.22E-308 |
| bmi | 0.719507 | <2.22E-308 | 0.719547 | <2.22E-308 | 0.719496 | <2.22E-308 |
| smk_stat | -0.88889 | 2.02E-84 | -0.89023 | 1.16E-84 | -0.88798 | 2.94E-84 |
| act_more | 0.75906 | 2.57E-77 | 0.758519 | 3.38E-77 | 0.759263 | 2.32E-77 |
| alc_less | -0.49658 | 4.92E-119 | -0.49147 | 9.31E-117 | -0.49821 | 8.73E-120 |
| salt_add | -0.81802 | 1.05E-118 | -0.81464 | 9.52E-118 | -0.81918 | 4.88E-119 |
| crp | 0.065327 | 1.40E-18 | 0.065339 | 1.39E-18 | 0.065363 | 1.34E-18 |
| apo_a | 7.25042 | <2.22E-308 | 7.25638 | <2.22E-308 | 7.248796 | <2.22E-308 |
| apo_b | 7.96638 | <2.22E-308 | 7.959616 | <2.22E-308 | 7.969412 | <2.22E-308 |
| edu | 0.029443 | 1.11E-50 | 0.0295 | 7.30E-51 | 0.029432 | 1.20E-50 |

We repeated the covariate-corrected analyses for systolic blood pressure, stratifying the analysis by (1) whether someone was on a BP medication, (2) age above or below 50, and (3) by race and ethnicity. We found that rs5051 C>T is associated with a 0.32 mmHg increase in systolic blood pressure in those not taking BP medications (p<0.001, n= 243,277), and a non-significant 0.08 mmHg increase in systolic blood pressure in those taking BP medications (p=0.399, n=67,725). The genotype groups followed the same non-significant pattern as the non-stratified analysis. Results were also not markedly different when stratified by age groups <50 or ≥50. The genotype group analysis results followed similar nonsignificant patterns in the race and ethnicity groups, however rs5051 C>T was associated with a 1.17 mmHg increase in systolic blood pressure in Black participants (p=0.032, n=4,580), a >4-fold larger effect size than seen in White participants (0.25 mmHg increase, p<0.001, n=294,582). **Figure 8A** shows the non-covariate adjusted numbers per rs5051 genotype group. Other race and ethnicity groups were not statistically significant. Interestingly, the rs5051 C>T variant has an 88% allele frequency in Black participants, compared to a 40% allele frequency in White participants in the UK Biobank data. **Figure 8B** shows phased haplotypes from the 1000 Genomes Project illustrating that the rs5051 T allele is far more common in those with African ancestry.

**Figure 8:**
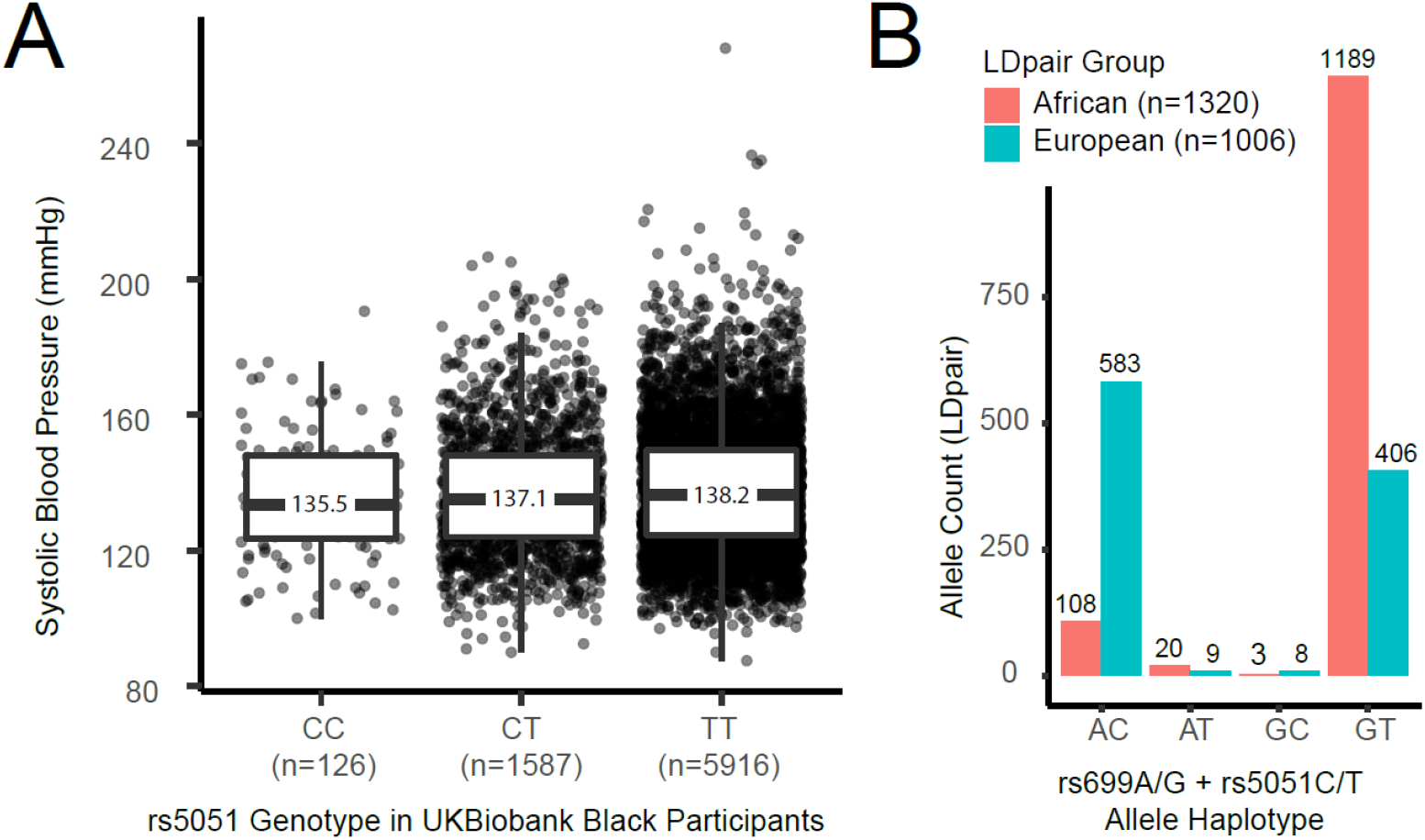
Analysis of (A) rs5051 C>T association with systolic blood pressure in Black participants in the UK Biobank data, and (B) allele haplotype frequencies for rs699 and rs5051 in the 1000 Genomes Project for those with African ancestry. Boxplot displays median (black bar), 25^th^ to 75^th^ percentile (box), and mean values are labeled over the box.

## DISCUSSION

In this study, we prioritized the strongest signal from our previous high-throughput variant screening assay (PASSPORT-seq) to further investigate the molecular mechanism and clinical relevance of rs699 A>G. We hypothesized that rs699 A>G was responsible for decreasing *AGT* abundance via increasing the binding strength to mir-122-5p, and, that when unopposed by the increased transcription caused by rs5051 C>T, would reduce *AGT* expression and be protective against HTN. Approximately 2-3% of people have an imbalance of these two SNPs, indicating the research question is of potential significance to the human population. Our *in silico* results support our hypothesis, but should be viewed conservatively since the change in binding strength was only around 3%. Our *in vitro* results (which isolate rs699 A>G) do not support our hypothesis, since we expected rs699 A>G to cause decreased *AGT* mRNA abundance in HepG2 hepatocytes (where mir-122-5p is highly expressed), but instead *AGT* mRNA was increased. In HT29 colon cells, rs699 A>G decreased *AGT* mRNA, demonstrating the potential for cell-type specific effects. Furthermore, the additional mir-eCLIP assay results confirm that rs699 is near a mir-122-5p binding site, but it is unlikely that the A>G variant strongly modifies mir-122-5p binding strength directly. *In vivo*, GTEx data did not show a strong liver upregulation of *AGT* in three individuals with unopposed rs5051 C>T, and did not show an obvious downregulation of *AGT* in three individuals with unopposed rs699 A>G. GTEx did demonstrate strong eQTLs with opposite direction of effects in different cell types, supporting that the functional effect of rs699 A>G or rs5051 C>T (since they are in LD) is cell-specific. Further analysis of retrospective clinical data from IU cancer patients did not show a clinically meaningful effect on several measures of HTN, though this analysis was likely underpowered and potentially confounded by cancer therapies. Retrospective UK Biobank analysis, which provided a large clinical population to isolate the independent effects of rs699 A>G and rs5051 C>T, also did not show a clinically meaningful effect on systolic blood pressure for the genotype group analysis based on our hypothesis. Taken together, our results suggest that rs699 A>G or rs5051 C>T has a cell-type specific functional role at the mRNA level, but its clinical significance on blood pressure and the development of HTN is small.

Our study has several limitations. The *in silico* analysis is taken out of the context of the secondary structure of the mRNA and does not account for the steric hindrance of RNA-binding proteins or N6-methyladenosine modifications that can alter the microRNA binding strength. Thus, we stress that the results of this analysis need to be interpreted conservatively. The *in vitro* results would have benefited from additional replicates conducted on different lots or different passages of cells with transfections occurring on separate days. Despite this limitation, however, our experiments were fit for the purpose of deciding whether to invest more into this line of research. The *in vivo* GTEx and clinical analyses in our institutional cohort were underpowered, but this limitation is inherent to retrospective data analysis where additional subjects cannot be recruited. Additionally, the tight linkage between rs699 A>G and rs5051 C>T necessitates the use of very large clinical cohorts to assess the independent effects of either SNP. In contrast, a great strength of our study is the combination of experiments and analyses spanning multiple disciplines of science (*in silico, in vitro, in vivo*, and observational) to test a single hypothesis. Another strength is that the UK Biobank analysis was overpowered and thus very useful in statistically rejecting the alternate hypothesis that rs699 A>G acts as a counteracting force against rs5051 C>T.

The initial observation that stimulated our work into rs699 was from our high-throughput assay testing the effect of 262 candidate SNPs on reporter mRNA abundance. While this assay was designed to screen for variants that modify microRNA binding, it does not do so directly. While this is a potential disadvantage, it is also a strength to identifying functional variants that exert their effect indirectly. Our results suggest that rs699 A>G may modulate the binding of mir-122-5p indirectly through other cell-type specific mechanisms such as altered recruitment of RNA-binding proteins or splicing machinery. In fact, ENCODE eCLIP experiments for RNA-binding proteins in HepG2 demonstrate that proteins involved in splicing and pre-mRNA processing; AQR, BUD13, CDC40, NOL12, PPIG, RBM15, RBM22, SRSF1, SRSF9, and TRA2A bind to *AGT* mRNA in the rs699 location, giving plausibility to this hypothesis (this data can be viewed at www.encodeproject.org).^48^ While not a main analysis of this paper, we used the GTEx liver *AGT* RNA-seq data to measure alternative splicing events at the exon junction near rs699 (3’ end of exon 2) to aid in interpreting these data.

We found that exon 2 was spliced to a non-canonical exon (i.e., not joined to canonical exon 3) on average 10% of the time compared to all exon 2 junction reads for each individual (range 0-21%, n=208). However, this alternative splicing did not correlate with rs699 genotype (nor did any single splice variant), indicating that rs699 does not interfere with *AGT* exon 2 splicing in the liver.

DROSHA, UCHL5, and XPO5 were also identified to bind in the rs699 location based on ENCODE data, but the strongest signal was for SND1 binding. SND1 is an endonuclease that mediates microRNA decay^49^ and is a component of the RNA-induced silencing complex (RISC),^50^ which is interesting given our finding that this site is near a mir-122-5p binding site. Thus, microRNA-mediated recruitment of SND1-containing RISC to this location is evident, but the clinical significance of this remains unknown.

A potentially important finding from this study is that rs5051 C>T and/or rs699 A>G had a significant association with systolic blood pressure in Black participants in the UK Biobank, demonstrating a 4-fold larger effect size than that seen in White participants. It is noteworthy that the allele frequencies for rs699 A>G and rs5051 C>T are approximately twice as common in African ancestry groups relative to Europeans. In addition to differences in minor allele frequency, it is known that the linkage disequilibrium blocks of *AGT* differ between White and Black people.^51^ Salt sensitive hypertensive phenotypes are known to be more common in individuals of African ancestry,^52-54^ which lends support to the possibility that rs5051 C>T or rs699 A>G mediate blood pressure changes more dramatically in Black patients through changes in *AGT* expression or abundance. Individuals with African Ancestry have lower plasma renin activity with preserved aldosterone levels,^55^ and substantial evidence also indicates that individuals of African ancestry respond better to calcium channel blockers and diuretics than beta-blockers or RAS acting antihypertensives.^56-58^ Aligning with this, the Eighth Joint National Committee (JNC 8) endorses different initial antihypertensive therapy for Caucasian and Black individuals. In individuals with normal kidney function, the JNC only recommends angiotensin converting enzyme inhibitor as first-line therapy in Caucasians.^59^ Here, we provide evidence that rs5051 C>T or rs699 A>G may be involved in hypertensive phenotype differences between individuals with European and African ancestry that may underly established differences in antihypertensive treatment efficacy between these groups. Despite this, significant admixture exists between populations. As access to whole genome sequencing in clinical care increases, studies like this will allow clinicians to move beyond the use of race as a surrogate for genotype.

In conclusion, the central findings from our study are that rs699 A>G is likely exerting an effect on *AGT* expression that is mediated by something which is cell-type specific, and that our hypothesis of opposing effects between rs699 A>G and rs5051 C>T can be statistically rejected. Further studies are warranted to determine if the reason for altered antihypertensive response in Black individuals might be due, in part, to the effect of rs5051 C>T or rs699 A>G.

## ACKNOWLEDGEMENTS

This work is supported by NIH-NIGMS T32GM008425 to NP, NIH-NIGMS Grant R35GM131812 to TCS, NIH-NIGMS Grant 1R01GM120156-01A1 to TL, NIH-NCI Grant 1R03CA223906-01 to TL, and NIH-NIGMS K23GM147805 to TAS.

The Genotype-Tissue Expression (GTEx) Project was supported by the [Common Fund](http://commonfund.nih.gov/GTEx/index) of the Office of the Director of the National Institutes of Health, and by NCI, NHGRI, NHLBI, NIDA, NIMH, and NINDS. The data used for the analyses described in this manuscript were obtained from the [GTEx Portal](http://www.gtexportal.org/) on 01/10/2023, and through dbGaP accession phs000424.v8.p2.

This research has been conducted using the UK Biobank Resource under Application Number 86441. Biospecimens were stored in the CTSI Specimen Storage Facility at Indiana University School of Medicine which is supported, in part, by grant NIH/NCRR RR020128. The content is solely the responsibility of the authors and does not necessarily represent the official views of the National Institutes of Health.

We thank all research participants involved in making this work possible.

## AUTHORS CONTRIBUTIONS

NRP designed, executed, and analyzed the experiments, and wrote the manuscript.

TS contributed to experimental design, analysis, and manuscript writing.

JL contributed to experimental execution.

MM contributed to experimental execution and manuscript writing.

RPK contributed to manuscript writing.

MTE, DL, TCS, and TL contributed to experimental design and manuscript writing.

## Notes

https://www.ncbi.nlm.nih.gov/geo/query/acc.cgi?acc=GSE227405

### Competing Interest Statement

The authors have declared no competing interest.

